# Antifungal activity of protamine

**DOI:** 10.1101/2024.12.07.627331

**Authors:** Sivakumar Jeyarajan, Anbarasu Kumarasamy

## Abstract

**Background/Objective:** Antimicrobial peptides (AMPs) are important innate defense molecules having wide spectrum of bioactivities like antibacterial, antifungal, antiparasitic and antiviral activities. The primary role of AMPs is to exert cytotoxicity on the invading pathogenic microorganisms and serve as immune modulators in higher organisms. Protamine is a polycationic peptide found in the nuclei of sperm in different vertebrate species has also antimicrobial activities. Protamine is thought to disrupt the cytoplasmic membrane caused by electrostatic interactions between the highly positively charged molecule and the negatively charged microbial cell surfaces. In addition to this, it is involved in wound healing activity and recruit leucocytes and modulators during inflammation. The present study proposes to investigate the anticandidal effects of protamine on clinical isolates of *Candida* spp. For experimental evaluation of anticandidal and antibiofilm activity, Minimal inhibitory concentration (MIC) was determined against *Candida albicans, Candida tropicalis* and *Candida krusei*. Protamine inhibited the growth of all the tested *Candida* spp. pathogens in respective MIC of 16 μg ml^-1^, 32 μg ml^-1^ and 256 μg mg^-1^. After MIC determination, the mechanism of action was evaluated by assessing structural and cellular biochemical changes. The structural changes were examined by scanning electron microscopy (SEM) after treating *C. albicans* with protamine and SEM images clearly showed membrane rupture indicating that the peptide targets the membrane. Biochemical changes like induction of reactive oxygen species (ROS) inside the cells which is essential for cell death was detected by staining the cells with 2′,7′-Dichlorofluorescin diacetate which turns to green colour in response to oxidative metabolism by esterification reaction. Protamine increased ROS production in C*andida* cells. These results suggested that protamine could be a lead compound for preparation of biomaterials for anticandidal treatment.

## 1. Introduction

Candidiasis, caused by *Candida* species, has become a significant global health issue in recent years [1, 2]. This is largely due to the growing resistance of these infections to existing antifungal medications like fluconazole, echinocandins, and polyenes. Vulvovaginal candidiasis (VVC) is a condition in women that causes vaginal discharge, pruritus, pelvic pain, and inflammation. Globally, 70–75% of women experience VVC at least once during their reproductive years [1, 2]. VVC can be acute or chronic, significantly impacting women’s quality of life by causing persistent discomfort, stress, and disruption to daily activities and mental health. The incidence of vaginitis caused by non-albicans *Candida* (NAC) species is increasing, posing significant challenges due to their reduced susceptibility to fluconazole, a commonly used drug. NAC infections are often recurrent.

VVC is frequently diagnosed based on non-specific symptoms and treated without culture confirmation in most of the developing and undeveloped countries. The widespread use of oral fluconazole for prophylaxis, especially in immunocompromised patients such as those with HIV [3] or organ transplant recipients, further contributes to resistance in *Candida* spp. [4]. Particularly NAC species, are inherently resistant to azoles which causes adverse reactions when administered to immunocompromised individuals. This alarms clinicians to explore for alternate treatment strategies.

Intrauterine devices (IUDs) are cost-effective and are widely used for women contraception. These devices are the sites of attraction for commensal microorganisms like bacteria and *Candida* spp. These commensals adhere to it and form biofilms. This biofilm can lead to a significant increase in their numbers, transforming the commensals into pathogens. IUDs can thus become reservoirs for *Candida* infections, accumulating larger biofilm masses and contributing to recurrent infections [5]. This pathogenicity causes inflammation in the surrounding tissues, leading to disease and a weakened immune system. The outgrowth of *Candida* spp. on these devices exacerbates vaginal candidiasis, causing alarm and loss of confidence among users. Thus, VVC can become invasive, leading to candidemia. The re-occurrence of infections may also lead to neoplasm causing epigenetic changes to tissues [6-10]. The progression from using a contraceptive device to a life-threatening condition is concerning and often overlooked. While amphotericin B is a last-resort antifungal treatment, it is generally not prescribed for vaginal candidiasis to avoid resistance development. Unfortunately, resistance to amphotericin B has also been observed. In cases of resistance, physicians may increase the dose, which is toxic to the patients and causes further complications. The World Health Organization (WHO) reports that immunocompromised IUD users are at higher risk of VVC [11]. The high mortality rate associated with these infections underscores the urgent need for effective infection control measures and anti-*Candida* treatments.

*Candida* spp. develops resistance by molecular mechanisms that are ultimately responsible for phenotype change in their cells. The molecular mechanisms involve over expression or mutation of target enzyme [12], alteration of the enzyme in the same biosynthetic pathway as target enzyme, upregulation of drug efflux pumps (ATP Binding Cassette), upregulation of target gene expression etc. The physical mechanisms include dimorphic transitions, i.e., phenotypic switching from a budding yeast cell to a filamentous form, which helps the pathogen evade the drug and adapt to invade host epithelial tissues [13]. Additionally, pleiotropic changes in target structures render the drug, incapable of binding to its target. The transformation into persister cells, which remain in a dormant state, makes molecular targets of *Candida* spp. inactive [14]. Furthermore, the formation of a matrix prevents anti-*Candida* drugs from reaching their targets. Most pathogenic *Candida* spp. are shielded by this matrix which facilitates biofilm formation. The biofilm formed on the matrix prevents antimicrobial substances to reach individual cells. Biofilm-associated cells require concentrations of fluconazole and amphotericin B that are 1000 times higher than those needed for planktonic cells [15, 16] to inhibit *Candida* biofilms. These characteristics plays a crucial role in the persistent recurrence of infections and antibiotic resistance These traits pose additional difficulties in orchestrating a successful treatment strategy for *Candida* infections, necessitating new therapeutics that can overcome the above problems.

Recent research is focused on cationic antimicrobial peptides (AMPs), which are toxic to *Candida* spp. but harmless to normal mammalian cells [17, 18]. This selectivity is due to the differences in membrane properties and composition between *Candida* spp. and mammalian cells. AMPs are composed of repeating sequences of positively charged and hydrophobic amino acids, giving them amphiphilic property. This amphiphilic property allows AMPs to bind to the negatively charged *Candida* spp. membrane (β-glucan, chitin, and phosphomannoproteins) and penetrate the hydrophobic membrane, forming channels that cause cytosolic content leakage. Since AMPs target negative charges rather than specific carbohydrate or lipid moieties, the likelihood of resistance is low. Histatins[19], protonectin [20], LL-37 in humans [21], N-terminal domain of bovine lactoferrin[22], epinecidin [23-26], synthetic helical peptides [27], and KABT-AMP [14] are few examples of AMPs having anti-*Candida* effect.

Protamine is a small, arginine-rich “MPRRRRSSSRPVRRRRRSRRRRRRGGRRRR” (NCBI accession # P69014) nuclear protein that replaces histone late in the haploid phase of spermatogenesis and essential for sperm head condensation and DNA stabilization. Protamine was discovered from Salmon fish testicles by Fredrich Miescher in 1869 and identified in the folding of nucleic acids in salmon sperm. Protamine has been approved by the FDA for injection or infusion into the blood vessel to treat heparin overdose [28]. It is also used as insulin carrier as isophane insulin and sold by Eli Lilly. Protamine has been used as gene delivery agent with minimal toxicity [29, 30]. Protamine has also shown to inhibit protease produced by bacteria [31]. Protamine has several characteristics, including high stability under heat, hence it is also used as a preservative in neutral or alkaline food [32].

In this study we experimentally determined the susceptibility of VVC isolates to protamine by microbroth dilution method to determine minimum inhibitory concentration (MIC), crystal violet staining, scanning electron microscopy (SEM) and reactive oxygen species (ROS) assay to study the mechanism of action.

## 2. Materials and Methods

### 2.1. Candida Strains, Peptides and Chemicals

Vulvovaginal candidiasis (VVC) isolates: *C. tropicalis* (strain CA4) and *C. krusei* (strain CA54) with high biofilm forming potential which were reported by Shanmugapriya et al. [33] were used to evaluate anti-*Candida* efficacy of protamine. The above two strains were present in major population in the isolates from women, aged 20–35 years who had signs of pelvic inflammation, hemorrhage and vaginal discharge etc. These two strains specially formed multiple layers of biofilm and contained several microcolonies. A pathogenic strain of *C. albicans* obtained from Microbial Type Culture Collection (MTCC 227) was used as control test organism.

### 2.2. Protamine and Chemicals

Protamine, Sabouraud Dextrose broth (SDB), Whatman sterile disks 2’,7’-dichlorofluorescein diacetate (DCFHDA) and glutaraldehyde were from Sigma Aldrich (Millipore Sigma Burlington, MA). All additional chemical reagents employed in the testing procedures were of analytical grade.

### 2.3. Assessment of Anti-Candidal Activity of Protamine Using Disk Diffusion

The anti-candida activity of protamine was evaluated by agar disk diffusion method. Briefly, one hundred microlitre of *Candida* spp. (10^6^ cells) were spread uniformly across Sabouraud dextrose agar culture plate individually. Protamine was dissolved in Phosphate buffer saline (PBS, pH 7.4) to 1 mg ml^-1^. Whatman paper disk impregnated with protamine at concentrations such as 10 μg, 20 μg, 30 μg, 40 μg and 50 μg were placed on the surface of the agar and incubated at 37°C for 24 hours so that the peptide diffuses from the filter paper into the agar. Post 24 hours, the zone of inhibition (ZOI) was measured and plotted as mean for three measurements,

### 2.4. Anti-Candidal Assay for Minimum Inhibitory Concentration (MIC) determination

Antimicrobial activity of protamine against *C. tropicalis* (strain CA4), *C. krusei* (strain CA54) and *C. albicans* (MTCC 227) were tested with the growth media: Sabouraud dextrose broth (SDB). Microbroth dilution method was used in accordance with the Clinical and Laboratory Standards Institute (CLSI) M27-Ed4 guidelines for anti-*Candida* susceptibility testing [34]. The stock solution of protamine was diluted with phosphate-buffered saline (PBS) to reach concentrations of 1, 2, 4, 8, 16, 32, 64, 128, 256 and 512 μg ml^-1^ [14]. Aliquots (20 μl) from each dilution were distributed to a 96-well polystyrene microtiter plate, and each well was inoculated with 180 μl suspension of *Candida* spp. in SDB containing 1× 10^6^ cells. Cultures were grown with gentle shaking for 24 h at 37 °C. Wells containing only cells without peptide was used as growth or positive control and plain broth was used as sterility or negative control. The absorbance was evaluated at 595 nm using a microplate reader (Bio-Rad, Hercules, CA, USA) and represented in terms of percentage of growth using growth control value as 100% growth. Minimal inhibitory concentration (MIC) of protamine was defined as the lowest concentration at which the percentage of growth less than 15% was observed. The antimicrobial assay was done in triplicates and the growth percentage was plotted as mean *±* S.D.

### 2.5. Biofilm Assay

To determine the ability of protamine to inhibit biofilm formation, 1 ml suspension of *Candida* spp. (1× 10^6^ cells) in SDB media were seeded into each well of a 24-well plate containing microscope coverslip (circular borosilicate cover glasses (15 mm), Fisher Scientific, Hampton, NH, USA) inserts [14]. The cover glass served as a substratum for microbial attachment. Protamine at its MIC concentration against *Candida* spp. were treated for 24 hours. For *C. albicans*: 32 μg ml^-1^, *C. tropicalis*: 64 μg ml^-1^ and *C. krusei*: μg ml^-1^ were used. After incubation, the spent media was aspirated and 1 ml of 0.1 % (w/v) crystal violet dissolved in PBS was added to each well and incubated for 30 min to stain the cells. Post staining, excess crystal violet was removed by washing twice with PBS. The cells were fixed with 4% formaldehyde, washed with PBS and dehydrated with 80% (v/v) ethanol. The stained cover slips were examined under light microscope with 40x magnification.

### 2.6. Scanning Electron Microscopy (SEM)

To view the morphological changes of *C. albicans* cells (MTCC 227) after treatment with protamine, SEM was employed [35]. In brief, *C. albicans* cells were grown to logarithmic phase with an inoculum size of (1× 10^6^ cells) on a microscope cover slip which was inserted to each well of a 24 well dish. Protamine at its MIC concentration of 32 μg ml against *C. albicans* was added to the culture medium. *C. albicans* grown without any peptide was used as negative control. After 6 h incubation at 37 °C, the cells were washed twice with phosphate buffered saline (PBS) pH 7.4, and metabolically fixed with an equal volume of 5% (v/v) glutaraldehyde at 4°C overnight. The metabolically fixed cells were dehydrated with serial gradient of ethanol wash from 70 to 100%. The coverslips were then sputter coated and examined under a scanning electron microscope (VEGA3 TESCAN, Czech Republic).

### 2.7. Measurement of Cellular ROS Production

ROS generated after treatment with protamine was measured by fluorometric assay with 2′,7′-dichlorofluorescin diacetate (DCFHDA) as described in [20]. Briefly, *Candida spp*. (1×10^6^ cells) were seeded into 24 well polystyrene plates and treated with or without peptide at their respective MIC for each species about 24 h. After 24 h treatment, the cells were incubated with 10 μM of DCFHDA for 1 h and washed with PBS pH 7.4. They were then visualized in fluorescent microscope (Accu-Scope, EXI-310, Commack, NY, USA) at 10x magnification and documented with green channel fluorescence intensities (excitation 488 nm and emission 525 nm respectively). Hoechst dye was used to stain the nucleus.

## 3. Results

### 3.1. Susceptibility of VVC Candida spp. isolates to protamine

The susceptibility of the VVC *Candida* spp. isolates to the antimicrobial peptide protamine was determined by agar disk diffusion method, also called zone of inhibition test or Kirby-Bauer method. It is a semi quantitative method to determine the ability of the peptide to inhibit *Candida* spp. growth or not. Protamine was added on the disk, and it diffuses through the agar to exert its function. If the peptide is effective against the *Candida* spp., no growth will be observed around the disk. This is the zone of inhibition (ZOI). The ZOI, evaluated for protamine against *Candida* spp. are shown in Figure 1 (a-c) and the measurements are shown in Figure 1 (d-f). As seen in the figure 1 a and b, *C. albicans* and *C. tropicalis* are sensitive to protamine however *C. krusei* is not sensitive to protamine at the concentrations tested (10 to 50 μg). A maximum ZOI of 1.5 cm was observed for *C. albicans* is 1.5 cm at 40 and 50 μg of protamine. For *C. tropicalis*, a gradient increase of ZOI was observed from 20 to 50 μg of protamine. However, for *C. krusei*, no ZOI was observed from 10 to 50 μg of protamine which made us to determine minimum inhibitory concentration (MIC) by broth dilution method where more quantity of protamine can be used for anti-*Candida* activity.

**Figure 1.**
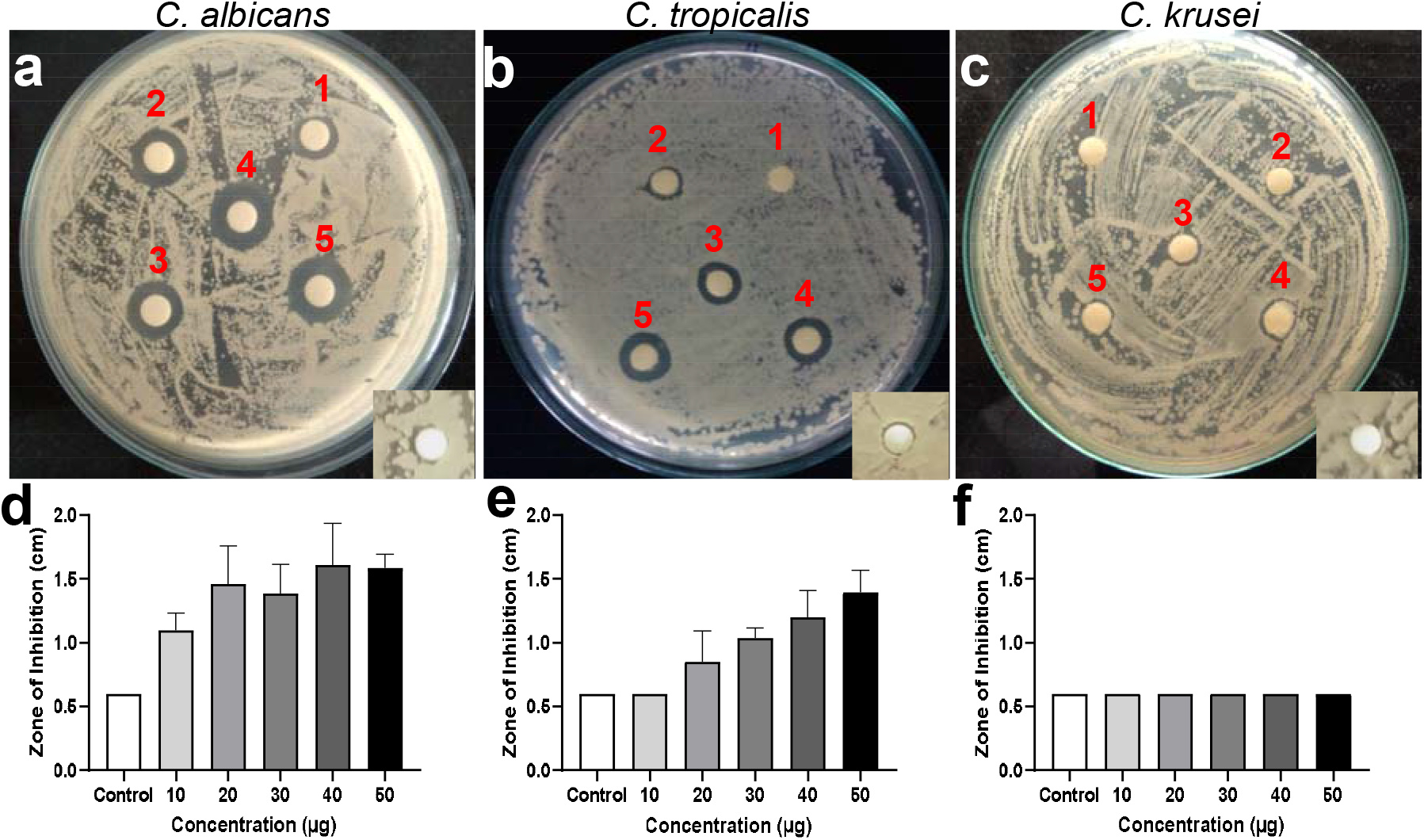
Susceptibility of a) *C. albicans* (MTCC 227), VVC *Candida* spp. isolates; b) *C. tropicalis* (CA4) and c) *C. krusei* (CA54) against protamine measured by agar diffusion assay. The numbers marked on the plate images describes the concentration of protamine loaded onto disks. (1-10, 2-20, 3-30, 4-40 and 5-50 μg). Inset shows the image of disk loaded with PBS as control. The bar graphs(d-f) shown below each image shows the diameter of zone of inhibition (ZOI) plotted for each concentration for each species. The control shows the diameter of disk alone (0.6 cm). For *C. albicans* and *C. krusei*, a concentration gradient increase in ZOI was observed. For C. *krusei*, no ZOI was observed for the measured concentration 10 to 50 μg

### 3.2. Protamine inhibit growth of VVC Candida spp. isolates

The *Candida*-cidal activity of protamine against the VVC isolates were studied using the microbroth dilution method for quantitatively determining their growth, as measured by optical density at 595 nm. Figure 2 shows the plot of growth of the isolates treated with protamine at the range of concentration (1 to 512 μg ml^-1^). The percentage of growth was measured by normalizing with the cells grown without peptides, which were considered 100%. The minimum inhibitory concentration (MIC) for protamine against the VVC isolates were determined from the lowest concentration at which the growth was inhibited. Protamine inhibited growth of *C. albicans* at 32 μg ml^-1^, *C. tropicalis* at 64 μg ml^-1^ and *C. krusei* at 256 μg ml^-1^. At this concentration, the wells inoculated with protamine showed no turbidity.

**Figure 2.**
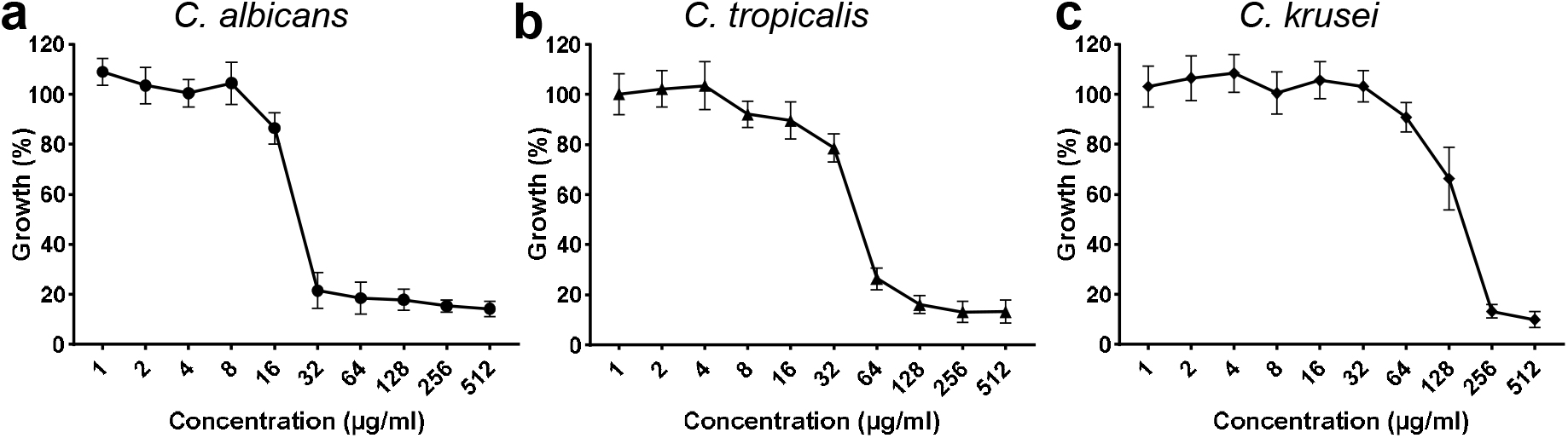
*Candida*-cidal activity of protamine against *Candida* spp. (a-c). Microbroth dilution assay was performed from 0 to 512 μg ml^-1^ to determine the MIC. Cell densities for *Candida* spp. measured at 595 nm without protamine are used as control with 100% growth The MIC of protamine against *C. albicans* is 32 μg ml^-1^, *C. tropicalis* is 64 μg ml^-1^ and *C. krusei* is 256 μg ml^-1^.

### 3.3. Protamine inhibit biofilm formation of VVC isolates

After determining the MIC, the ability of protamine to prevent the biofilm formation of VVC isolates formed on glass cover slips was determined by crystal violet staining. As these isolates were reported to form biofilm [33], they were cultured with and without protamine at its MIC against the VVC isolates for 24 h. After the treatment period, the coverslips containing the *Candida* spp. cells were stained with crystal violet, and then examined under an inverted 40x light microscope Figure 3. The images shown in Figure 3 (d -f), provide visible evidence of the protamine’s ability to inhibit biofilm formation. Control samples (*Candida* spp. cells without peptides) showed dense cell clusters indicative of substantial biofilm. The protamine treated cover glass did not have dense biofilm matrix and showed a significant reduction in cell numbers

**Figure 3.**
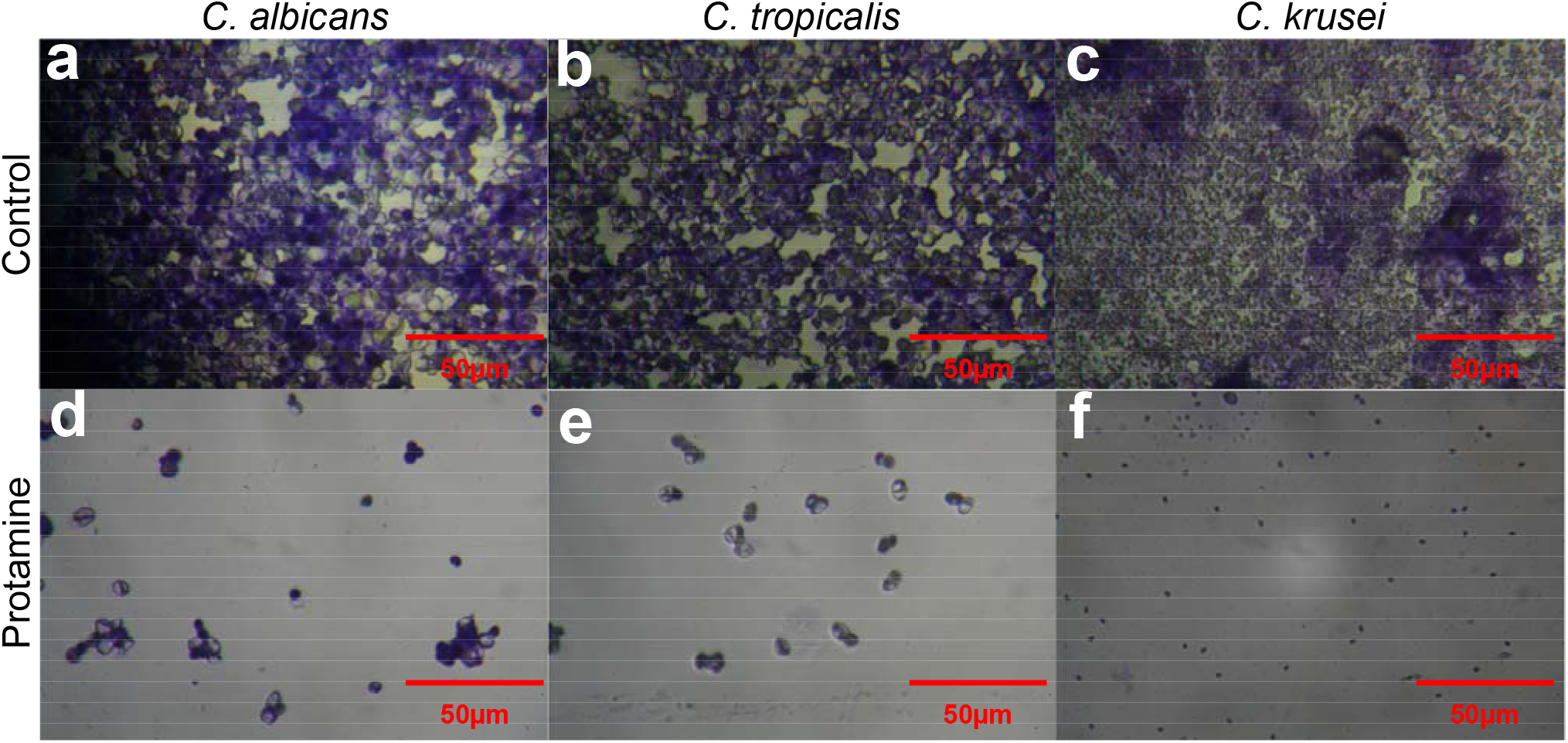
Light microscopic image of crystal violet stained control cells a) *C. albicans*, b) *C. tropicalis* (CA4) and c) *C. Krusei* (CA54) at 40x magnification. protamine (d-f) were treated with 32, 64 and 256 μg ml^-1^ of protamine respectively for 24 hours and stained with crystal violet.

### 3.4. Protamine disrupt C. albicans Membrane integrity

Scanning Electron Microscopy (SEM) was utilized to investigate the impact of protamine on membrane damage and penetration efficiency in the *C. albicans* cells as illustrated in Figure 4. Under SEM, the untreated control *C. albicans* cells are seen as a layer of cells as a biofilm and exhibited typical features: oval shapes, smooth surfaces, polar buds, and bud scars. The protamine treated *C. albicans* cells (32 μg ml^-1^ for 6 h) are less in numbers in the field and there are spaces between the cells. The cells have undergone plasmolysis, causing shrinkage. Some cells exhibit surface deformations, including rough textures, disruption, structural destabilization and cell wall collapse.

**Figure 4.**
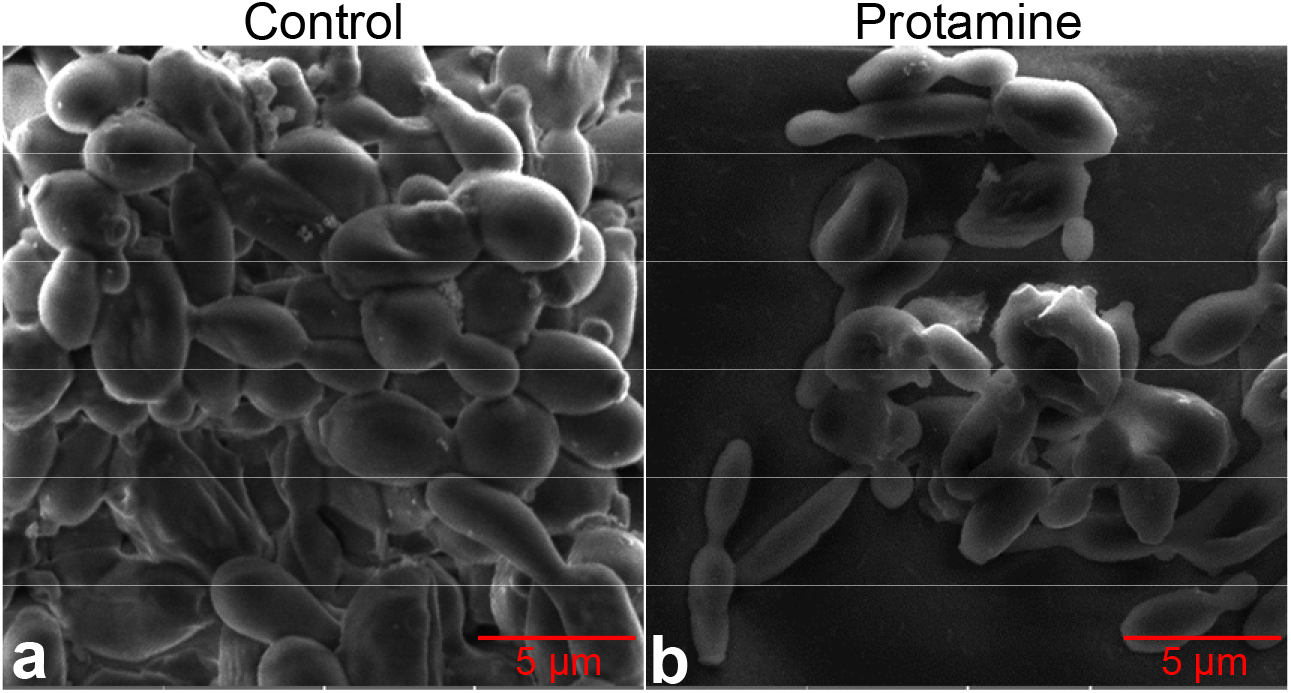
Scanning electron microscopic (SEM) image of *C. albicans* (MTCC 227) (a) Control, and (b) protamine treated cells. *C. albicans* was treated with protamine at 32 μg ml^-1^ for 6 h. The reduction in the number of cells noted as empty spaces between the cells and the disruption of cell membrane demonstrates the membrane disruption activity of protamine. The protamine treated cells show plasmolysis and ruptured surface with internally collapsed morphology due to cytoplasm leakage and appears shrunk with grooves compared to control cells.

### 3.4. Protamine treatment induce ROS generation

The present study evaluated whether the membrane perturbation generated ROS because of more complex intracellular signaling events, like oxidation of phospholipids and other macromolecules. Protamine’s ability to induce ROS formation was analyzed by using the cell permeant fluorogenic dye DCFDA. DCFDA is a cell-permeant dye that is oxidized to yield fluorescence when exposed to ROS. The fluorescence can be monitored with the excitation wavelength of 488 nm and the emission wavelength of 525 nm. As illustrated in Figure 5, no green fluorescence is observed in control *Candida* spp. The phase image shows dense cell population. Protamine treatment showed reduced cell density by phase and induced ROS production, inferred by the presence of green colored cells in the field.

**Figure 5.**
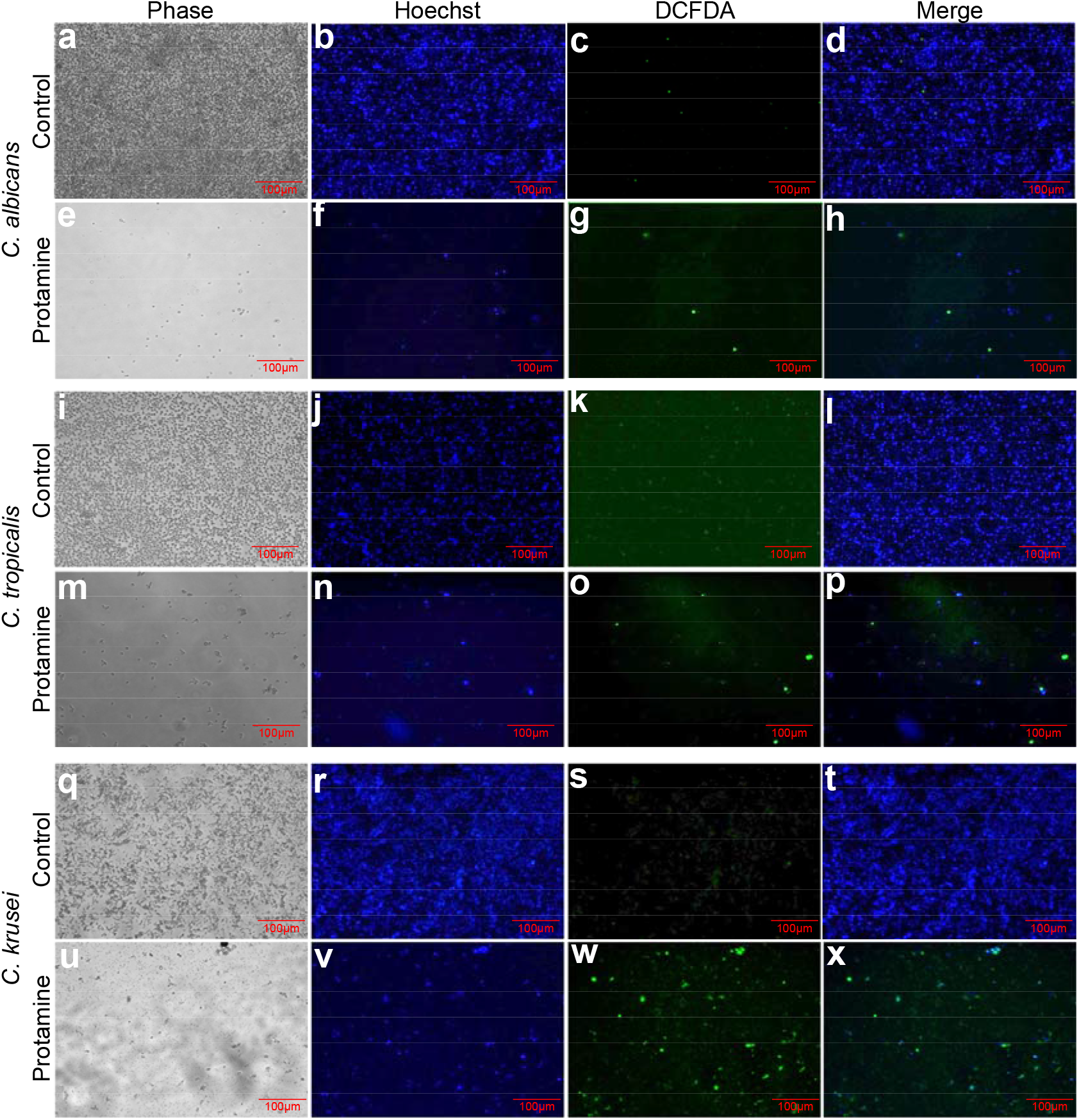
Effect of protamine on ROS production in *Candida* spp. Protamine at its MIC was added to VVC *Candida* spp. isolates, and ROS generation was studied. ROS generation was studied using DCFHDA pre-incubated with a 20 μM concentration in *Candida* spp. grown for 24 hours at 37°C. After 24 hours, the culture dish was washed with PBS (pH 7.4) and observed under a fluorescent microscope. The images were captured at an excitation wavelength of 488 nm and an emission wavelength of 525 nm. Top two panels show *C. albicans* without (a-d) and with (e-h) protamine. Couple of rows in the middle panels represents *C. tropicalis* treated with (m-p) and without (i-l) protamine. C. *krusei* with (u-x) and without (q-t) protamine is shown in the bottom two panels. Control represents cells alone and protamine panel represents cells treated at their MIC.

## 4. Discussion

According to the center for disease control (CDC), candidemia accounts for 25% of mortality among the patients hospitalized due to implanted device-related bloodstream infections [36] [37]. *C. albicans* has been the most prominent species causing Candidiasis, however, in recent years there are more occurrences of increasing population of non-albicans *Candida* (NAC). General prescription of azoles to treat Candidiasis is the cause of NAC increase, because NAC are inherently resistant to fluconazole, a commonly prescribed drug for Candidiasis. This is the primary reason for evaluating the susceptibility of VVC isolates; *C. tropicalis* and *C. krusei* to protamine in this study.

Antimicrobial peptides (AMPs) are short cationic amphiphilic peptides which exhibit helical or beta-sheet structures that facilitates the peptide to penetrate the microbial cells specifically because of their anionic membrane (β-glucan, chitin, and phosphomannoproteins) [38] [39]. The short AMPs also form aggregates that will cause cell death [14, 23, 27, 40-44] as defined by carpet model mechanism of action [45]. AMPs have been studied pre-clinically and clinically to treat Candidiasis. Human lactoferrin-derived peptide hLF1-11 was studied in clinical trials for anti-Candidemia effect [46, 47]. Synthetic AMP; PL-18 is under clinical trial for treating vaginal colpomycosis, bacterial vaginosis, and mixed vaginitis. PL-18 is loaded into suppository for this purpose [48]. CZEN-002, a synthetic octapeptide, derived from α-melanocyte stimulating hormone (α-MSH) was studied for treating vulvovaginal candidiasis as topical application.

Fascinated by the potential of AMPs to specifically kill pathogens, we evaluated the anti-biofilm potential of an arginine-rich AMP named protamine. Protamine has been reported to have antimicrobial activity against periodontopathic bacteria such as *Porphyromonas gingivalis, Prevotella intermedia* and *Aggregatibacter actinomycetemcomitans* [49]. Protamine has also been reported to show antimicrobial activity against catheter-associated *E. coli, Klebsiella pneumoniae, Pseudomonas aeruginosa, Staphylococcus epidermidis*, and *Enterococcus faecalis* biofilms in combination with ciprofloxacin [50] or chlorhexidine [51]. It also acts as an N-acetyl-D-glucosamine-1-phosphate acetyltransferase (GlmU) inhibitor to inhibit cell wall biosynthesis in Gram-negative and Gram-positive bacteria [52]. Additionally, it has been shown to possess anti-Candidal activity against *Candida albicans* isolated from poly (methyl methacrylate) (PMMA) dental prostheses [53].

Because protamine was reported to have antimicrobial activity against urinary catheter and dental prostheses-associated microbes, we evaluated its anti-Candidal and anti-biofilm activity against IUD-associated VVC non-albicans Candida isolates. The results from *in-vitro* evaluation showed that the MIC of protamine is 32, 64, and 256 μg ml^-1^ against *C. albicans, C. tropicalis*, and *C. krusei*, respectively. At these MICs, protamine exhibited significant level of anti-biofilm activity. SEM analysis of *C. albicans* clearly demonstrated that protamine binds to the membrane and disrupts its organization. This disruption is seen as a roughened membrane surface and the presence of pores. These results correspond well with previously reported mechanisms, such as the dilation of ionic channels by protamine sulfate, which facilitates the transport of antibiotics to the cytoplasm [54]. The mode of action for protamine is also thought to involve the disruption of the cytoplasmic membrane caused by electrostatic interactions between the highly positively charged peptide and the negatively charged cell surface. This electrostatic interaction mediates disruption, resulting in the leakage of K^+^, ATP, and intracellular enzymes from the cells treated with protamine [44, 55, 56]. Based on the SEM results obtained, we can compare the mechanism of action reported for protamine studied on bacterial membranes. Protamine interacts with the phospholipids of the bacterial cell membrane, increases membrane-ATPase activity, increases membrane permeability, and facilitating the delivery of antimicrobial compounds to the cytoplasm [57]. For *Candida* cells, we can compare the phosphomannoproteins to phospholipids. Furthermore, protamine generated ROS in the studied *Candida* spp. This may be due to membrane disruption, which in turn triggers the production of ROS within the cells like other AMPs previously reported from plant defensin [58]. Since the peptides form pores, this will lead to exposure to the oxygenic environment which increases the ROS to the damage the membrane. However, the series of events needs to be studied further in a series of steps.

Since protamine is clinically used to counteract heparins preoperatively or during cardiac surgery, it also has potential applications as an anti-candidal agent to disrupt biofilm on IUD. The median lethal dose (LD50) of protamine sulfate reported for rabbits and mice was found to be between 200 mg/kg and 300 mg/kg when given subcutaneously [51]. This gives evidence that coating IUD with protamine will be highly beneficial to inhibit the biofilm formation and does not cause harm to host.

## 5. Conclusion

Protamine demonstrated potency in destroying membrane integrity and preventing biofilm formation of VVC isolates. These quantitative attributes prove the potential application of protamine as anti-biofilm agent on medical implants.

## Funding and Acknowledgements

This research was funded by the Department of Biotechnology, India, (**Ref. No**. BT/PR2071/BBE/117/241/2016). RUSA 2.0 Biological Sciences (**Ref. No**. 311/RUSA(2.0)/2018), Bharathidasan University, Tiruchirappalli and Tamil Nadu State Council for Higher Education (TANSCHE) (**Ref. No**. 01706/P6/2021). S.J acknowledges Indian Council of Medical Research (ICMR-NET61754/2010) for granting fellowship. We thank National College Tiruchirappalli, India for performing Scanning Electron microscopy studies.

## Abbreviations

ABC: ATP Binding Cassette
AMP: Anti microbial Peptide
CDC: Centre for Disease Control
DCFHDA: 2′,7′-dichlorofluorescein diacetate
MIC: Minimum Inhibitory Concentration
PBS: Phosphate Buffered Saline
ROS: Reactive Oxygen Species

